# Reproducibility of PD patient-specific midbrain organoid data for in vitro disease modelling

**DOI:** 10.1101/2025.04.17.649331

**Authors:** Elisa Zuccoli, Haya Al Sawaf, Mona Tuzza, Sarah Nickels, Alise Zagare, Jens C. Schwamborn

## Abstract

Midbrain organoids are advanced in vitro cellular models for disease modelling. They have been used successfully over the past decade for Parkinson’s disease (PD) research and drug development. The three-dimensional structure and multicellular composition allow disease research under more physiological conditions than is possible with conventional 2D cellular models. However, there are concerns in the field regarding the organoid batch-to-batch variability and thus the reproducibility of the results. In this manuscript, we generate multiple independent midbrain organoid batches derived from healthy individuals or GBA-N370S mutation-carrying PD patients to evaluate the reproducibility of the GBA-N370S mutation-associated PD transcriptomic and metabolic signature as well as selected protein abundance. Our analysis shows that GBA-PD-associated phenotypes are reproducible across organoid generation batches and time points. This proves that midbrain organoids are not only suitable for PD *in vitro* modelling, but also represent robust and highly reproducible cellular models.

## Introduction

Parkinson’s disease (PD) is a common neurodegenerative disorder characterized by the progressive loss of midbrain dopaminergic neurons, leading to motor symptoms such as bradykinesia, rigidity, and tremors, as well as cognitive impairments (Kalia & Lang, 2015). Conventional two-dimensional (2D) cell cultures and animal models have limitations in replicating the complexity of the human midbrain, hindering their utility in understanding PD pathogenesis, and developing effective therapies (Kelava & Lancaster, 2016). Specifically, 2D cultures lack intricate cell-to-cell interactions, which are present in three-dimensional structures, and are crucial for accurately modelling neural environments. Additionally, animal models, such as mice, do not naturally develop neurodegenerative diseases like Parkinson’s, and their brain anatomy differs significantly from humans which limits their effectiveness in studying complex human brain disease processes (Dovonou et al., 2023). Recent breakthroughs in stem cell technologies have resulted in the development of three-dimensional (3D) midbrain organoids, which more closely mimic the architecture and cellular diversity of the human midbrain. Importantly, these organoids are derived from patient-specific induced pluripotent stem cells (iPSCs) and are generated via the differentiation of iPSCs into neuroepithelial stem cells (NESCs), which subsequently self-organize into midbrain-like structures (Monzel et al., 2017; Nickels et al., 2020). This approach offers a physiologically relevant platform to study disease mechanisms and to test therapeutic interventions on patient-specific genetic backgrounds. GBA-associated PD, linked to mutations in the glucocerebrosidase (GBA1) gene, represents one of the most significant genetic risk factors for PD (Smith & Schapira, 2022). GBA mutations impair lysosomal function and autophagy, contributing to α-synuclein protein accumulation and dopaminergic neuron loss (Smith & Schapira, 2022). Additionally, cellular senescence has emerged as a contributing factor in PD, with evidence suggesting that senescent cells exacerbate neurodegeneration through pro-inflammatory pathways (Chinta et al., 2018; Melo dos Santos et al., 2024; Wang et al., 2024). Midbrain organoids derived from GBA-PD patients have shown hallmarks of disease, including dopaminergic neuron loss, oxidative stress, impaired lipid metabolism, and a senescence-associated phenotype (Rosety et al., 2023). These findings underline the value of midbrain organoids as a robust platform to explore the interplay between genetic risk factors, cellular senescence, and neurodegeneration. iPSC-derived organoid models hold the potential to inform therapeutic strategies targeting the underlying mechanisms of GBA-associated PD and related neurodegenerative disorders.

Although midbrain organoid models hold significant promise, they also present certain limitations that must be carefully considered when designing experiments. Complex, multistep culturing protocols can increase sources of variability. Thus, increasing the number of replicates needed both within a batch and between independent batches. Therefore, it is important to be aware of variation sources in order to account for it. In this manuscript, we explore these sources of variation and inform on the relevance of cell passage in reducing batch-to-batch variability. Overall, our results show that key disease phenotypes are reproducible despite potential sources of variability.

## Materials and Methods

### Ethical approval

The work with iPSCs has been approved by the Ethics Review Panel (ERP) of the University of Luxembourg and the national Luxembourgish Research Ethics Committee (CNER, Comité National d’Ethique de Recherche) under CNER No. 201901/01 (ivPD) and No. 202406/03 (AdvanceOrg).

### Midbrain organoid culture

The patient GBA1 cell line was obtained from the European Bank for induced pluripotent stem cells (EBiSC), the patient GBA2 cell line was provided by the University College London and the patient GBA3 cell line from the Coriell Institute. The healthy control 1 and 2 were generated at the Max Planck Institute and healthy control 3 was provided by the Coriell Institute. One iPSC line from each donor was used, which was derived into NESCs as described in Reinhardt et al (2013). Midbrain organoids were generated from NESCs according to a protocol previously published by Monzel et al and Nickels et al (2017, 2020) until day 30 or day 60 of organoid culture depending on the experiment. NESCs were cultured in N2B27 maintenance media in 6-well geltrex (Thermo Fisher Scientific, cat.no. A1413302) precoated plates. The N2B27 base media consists of DMEM-F12 (Thermo Fisher Scientific, cat.no. 21331046) and Neurobasal (Thermo Fisher Scientific, cat.no. 10888022) in a 50:50 ratio and is supplemented with 1 % GlutaMAX (Thermo Fisher Scientific, cat.no. 35050061), 1 % penicillin/streptomycin (Thermo Fisher Scientific, cat.no. 15140122), 1% B27 supplement w/o Vitamin A (Life Technologies, cat.no. 12587001) and 2% N2 supplement (Thermo Fisher Scientific, cat.no. 17502001). For the maintenance of NESCs, the N2B27 base media was supplemented with 150 µM ascorbic acid (Sigma, cat.no. A4544), 3 µM CHIR-99021 (Axon Medchem, cat.no. CT 99021), and 0.75 µM purmorphamine (Enzo Life Science, cat.no. ALX-420-045). The NESC maintenance media was exchanged every second day. For the generation of midbrain organoids, NESCs were detached at 80% confluence with Accutase (Sigma, cat.no. A6964). Trypan Blue was used to count the number of viable cells. 9,000 live cells were seeded into each well of a 96-well ultra-low attachment plate (faCellitate, cat.no. F202003) in 150 µl of NESC maintenance media to initiate spheroid formation, marking day 0 of organoid culture. On the 2^nd^ day of midbrain organoid culture, the maintenance media was changed to patterning media, where N2B27 base media was supplemented with 200 µM ascorbic acid (Sigma, cat.no. A4544-100G), 500 µM dbcAMP (STEMCELL Technologies, cat.no. 100-0244), 10 ng/ml hBDNF (Peprotech, cat.no. 450-02-1mg), 10 ng/ml hGDNF (Peprotech, cat.no. 450-10-1mg), 1 ng/ml TGF-β3 (Peprotech, cat.no. 100-36E) and 1 µM purmorphamine (Enzo Life Science, cat.no. ALX-420-045). The next media change with the pattering media was done on the 5^th^ day of organoid culture. On the 8^th^ day of organoid culture, the patterning media was replaced by the differentiation media, which excluded PMA from the patterning media composition. Further media changes were done every 3-4 days until sample collection. NESC and midbrain organoid cultures were regularly (once per month) tested for mycoplasma contamination using LookOut® Mycoplasma PCR Detection Kit (Sigma, cat.no. MP0035-1KT).

### Whole mount staining

Midbrain organoids were collected at day 30, fixed with 4% paraformaldehyde (PFA) overnight at 4 °C and washed 3x with PBS for 10 min. The whole organoids were permeabilized and blocked with 1% Triton X-100 and 10% donkey serum in PBS for 24 h at room temperature (RT) and washed with 0.01% Triton X-100 in PBS for 30 min at RT on an orbital shaker. The organoids were incubated for 4 days at 4 °C on an orbital shaker with the primary antibodies (Supplementary Table 2) diluted in 0.5% Triton X-100 and 3% donkey serum in PBS. They were washed 3x with PBS for 1 h before incubation for 2 days at 4 °C on an orbital shaker with secondary antibodies (Supplementary Table 2) diluted in 0.5% Triton X-100 and 3% donkey serum in PBS. The whole organoids were washed 3x with 0.05% Triton X-100 in PBS and once with Milli-Q water for 5 min at RT before mounting them with Fluoromount-G mounting medium. (Southern Biotech).

### Immunofluorescence staining of midbrain organoid sections

Midbrain organoids were collected at day 30, fixed with 4% paraformaldehyde (PFA) overnight at 4 °C and washed 3x with PBS for 10 min. Four midbrain organoids per cell line were embedded in 3% low-melting point agarose (Biozym, 840100) and the solid agarose block with the assembloid was sectioned at 60 μm using the vibrating blade microtome (Leica VT1000s, RRID:SCR_016495). The sections were permeabilized for 30 min in 0.5% Triton X-100 at RT, followed by one quick wash with 0.01% Triton X-100 in PBS. The sections were then blocked for 2h at RT with blocking buffer containing 2.5% BSA, 2.5% donkey serum, 0.01% Triton X-100 and 0.1% sodium azide in PBS. Primary antibody (Supplementary Table 2) was diluted in blocking buffer and the sections were incubated with the primary antibody dilutions for 48h at 4°C on an orbital shaker. Incubation with secondary antibodies and mounting of the sections were performed as described by Nickels et al. (2020).

### Image 3D reconstruction

The 3D structure of the dopaminergic neuron was reconstructed from whole mount stainings using IMARIS software (version 9.9.1) (Bitplane, RRID:SCR_007370). Z-stack planes were used to visualise the fragmentation of dopaminergic neurons positive to tyrosine hydroxylase (TH) in GBA-PD compared to hearty control (WT) midbrain organoids.

### β-galactosidase staining

60 μm sections from midbrain organoids were used in the β-galactosidase staining, using the Senescence Detection Kit (Abcam, cat.no. ab65351). Two sections from one midbrain organoid per cell line were used. Images were acquired at 4x and 10x on an Olympus IX83 Automated Fluorescence Microscope (RRID:SCR_020344) for qualitative images and enhanced using Fiji software (RRID:SCR_002285) to account solely for differences in the background levels of light.

### Image acquisition and analysis

For high-content imaging, mounted organoids were scanned using the Yokogawa CellVoyager CV8000 microscope (RRID:SCR_023270). A 4x pre-scan in the 405 channel identified organoid-containing wells, enabling the creation of masks to outline organoids. These masks guided the selection of the field for imaging at different wavelengths with a 20x objective. For all stainings, three to four organoids per condition and four batches were analyzed, with details provided in figure legends. Qualitative images were captured using a Zeiss LSM 710 Confocal Inverted Microscope (RRID:SCR_018063) with 20x, or 60x objectives.

Immunofluorescence images of the whole mount organoids from the Yokogawa microscope were analyzed in MATLAB (2021a, Mathworks, RRID:SCR_001622) using a custom image-analysis algorithm as described by Bolognin et al (2018). The algorithm merges overlapping sections into mosaic images, smoothes and combines color channels and removes small objects. Masks were created and refined for each marker based on pixel intensity to quantify marker areas in 3D space (voxels). Statistical significance was calculated using Wilcoxon T-test in R. Statistically significant results were indicated when p values were * < 0.05, ** < 0.01, *** < 0.001 and **** < 0.0001.

### RNA extraction, library preparation and sequencing

Total RNA was extracted from each organoid generation batch, with 15 to 20 midbrain organoids pooled per batch at either day 30 or day 60. RNA isolation was done using the RNeasy Mini Kit (Qiagen, cat.no. 74106) following the manufacturer’s protocol. Messenger RNA was purified from total RNA using poly-T oligo-attached magnetic beads. After fragmentation, the first strand cDNA was synthesized using random hexamer primers, followed by the second strand cDNA synthesis using either dUTP for directional library or dTTP for non-directional library. Library preparations were sequenced on an Illumina platform by Novogene’s sequencing service.

### Transcriptomic analysis

RNA sequencing data were pre-processed on the Galaxy server version 23.2.rc1 following Galaxy training tutorial (Hiltemann et al., 2023, Doyle et al., 2024). Reads were mapped to the human reference genome hg38 using the HISAT2 tool. Mapped reads were counted using the featureCounts function on the BAM output files of HISAT2. Differential expression analysis was conducted in the R statistical programming software version 4.4.2. (RRID:SCR_001905) using the software package “DESeq2” (version 1.42.1, RRID:SCR_015687) (Love et al., 2014). P-value significance scores for differential expression were adjusted for multiple hypothesis testing according to the Benjamini and Hochberg method (Benjamini & Hochberg, 2005). Pathway enrichment analysis was performed with the Enrich R tool (Kuleshov et al., 2016) using the gene-level differential expression results table obtained with the DESeq2. We made use of two Gene-set libraries: the “Reactome Pathways 2024” library (Milacic et al., 2024) and the “GO Biological process 2023” library (Ashburner et al., 2000, Aleksander et al., 2023). Euclidean distance analysis was performed in R using base functions for distance computation and the “dplyr” package (version 1.1.4, RRID:SCR_016708) for data manipulation, ensuring precise distance measurements through established statistical functions (Shirkhorshidi et al., 2014, Kim et al., 2019). This approach allowed us to quantify the relative positioning of the data points within the multidimensional space defined by the clusters, providing insight into its similarity to each group.

### Analysis of explained variance

To assess the sources of variability in gene expression data, Principal Variance Component Analysis (PVCA) was performed in R using the package version 1.42.0 (Bushel et al., 2024). PVCA performs the dual analysis Principal Component Analysis (PCA) and Variance Component Analysis (VCA). This results in a reduction in data dimensionality while retaining most of its variability. A mixed linear model is then applied, treating all factors of interest as random effects, including two-way interaction terms, to estimate and partition the total variance attributed to each term. This method accounts for all defined sources of variability, including both experimental (cell line, passage, batch) and biological (disease status, patient sex) factors. Both, biological and experimental variates were treated as random factors and the variance attributed to other technical variables was assigned to a residual variance category.

### Transcriptomics and metabolomics integration

RNA sequencing and metabolomics data were integrated using MixOmics package (version 6.22.0, RRID:SCR_016889) (Rohart et al., 2017) in R (version 4.2.2) with DIABLO framework (Singh et al., 2019). The separation between sample groups was assessed using the correlation of the top 50 genes with the top five metabolites from the first component of Partial Least Squares Discriminant Analysis (PLS-DA). Pearson correlation (abs(r) = 0.85) between metabolomics and transcriptomics features was further visualized in a Circos plot (RRID:SCR_011798).

### Sample size estimation using power analysis

Analysis was done in R programming language (version 4.2.2). For the RNAseq dataset statistical power based on sample size was estimated using PROPER package (version 1.30.0). Library size dispersion and baseline expression values were determined based on the complete RNAseq data (dataset 1) of midbrain organoid samples from eight independent organoid batches with 48 samples per group (3 cell lines x 2 independent groups (WT and GBA-PD) x 8 independent organoid batches). 20 simulations were run using “DESeq2” (version 1.42.1) (Love et al., 2014) as a differential expression analysis method for possible sample sizes considering that we have three independent cell lines per group (WT or GBA-PD), thus sample size can increase by three for each batch (3 cell lines x 2 batches = 6 samples per group). To determine the relevant effect sizes between the two groups, we estimated the median, min and max log2 fold change of significantly differentially expressed genes between GBA and control sample groups. The false discovery rate was set to 5%.

For metabolomics and imaging experiments, sample size estimation was done using pwr package (version 1.3.0) with a defined significance of alpha=0.05 and power=0.8. For both datasets Cohen’s effect size was calculated using the effsize package (version 0.8.1) for significantly different abundant metabolites and TH normalised to nuclei of imaging features respectively. Based on normal data distribution of metabolomics data, we applied pwr.t.test function, while considering potential unequal sample sizes in each group and non-normal data distribution of imaging data, function pwr.2p.test of the pwr package was applied to imaging data.

## Results

### Disease and cell passage are key sources of variation in midbrain organoid culture

The midbrain organoid culture system is influenced by several factors, including cell line selection, initial passage number, donor sex, and potential batch effects from different experimental runs, all of which require careful consideration (Figure 1, Table 1). To investigate each factor’s contribution to the data variance we generated midbrain organoids from three healthy donors and three PD patients carrying the GBA-N370S variant (GBA-PD) (Supplementary Table 1). iPSCs derived from GBA-PD patients and healthy donors were differentiated into neuroepithelial stem cells (NESCs) and patterned towards a midbrain identity to generate midbrain organoids (Monzel et al., 2017). In this study, two independent datasets were produced, where Dataset 1 comprised RNA sequencing (RNAseq) and immunofluorescence analyses performed on 30-day-old midbrain organoids, and Dataset 2 included metabolomic analysis and RNAseq analysis performed on both day 30 and day 60 midbrain organoids cultures. While Dataset 1 was used to define experimental and biological sources of variation, Dataset 2 served to validate the reproducibility of key phenotypes across independent experiments (Figure 1).

**Figure 1.**
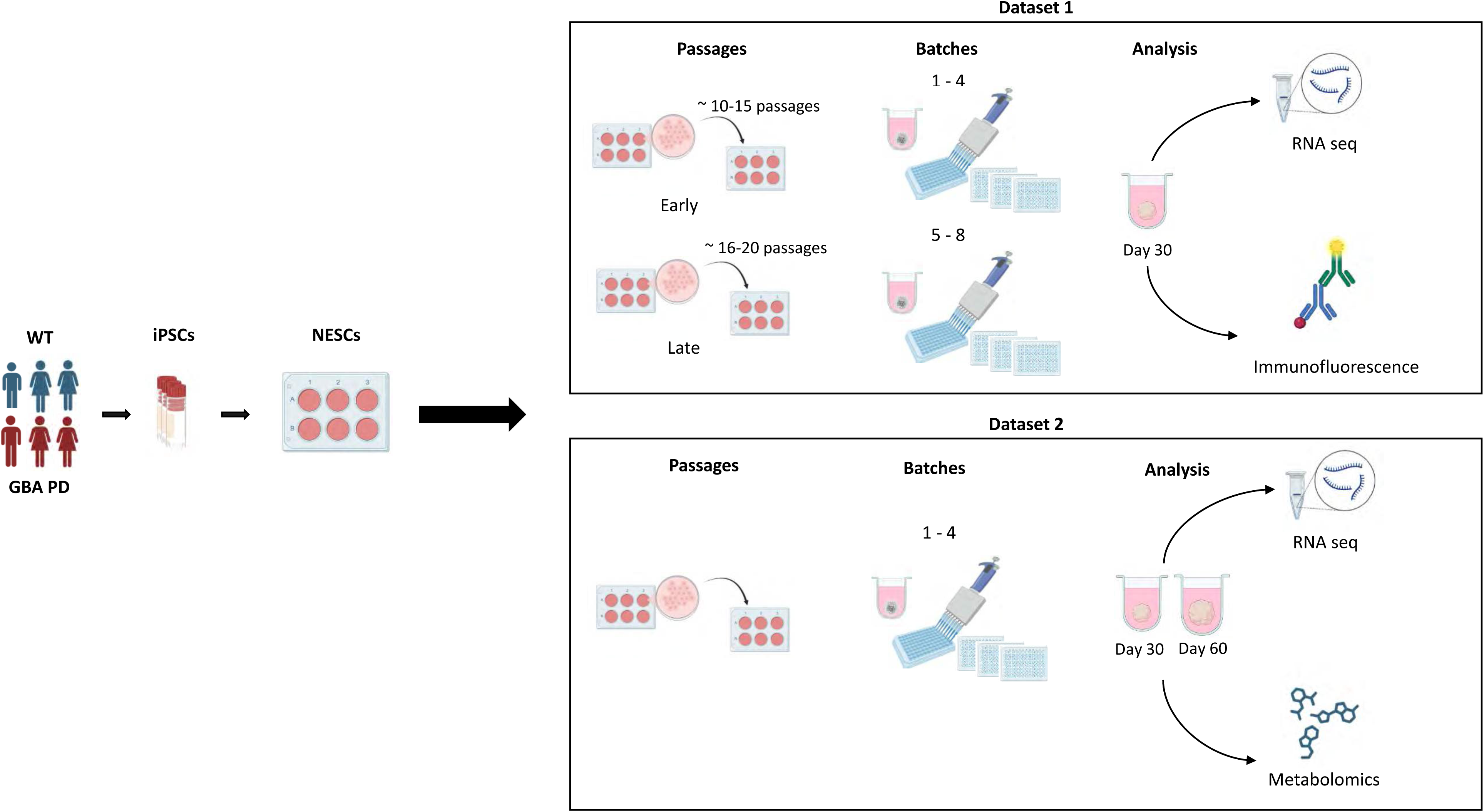
Overview of Factors Affecting Midbrain Organoid Culture and Dataset Generation. Schematic overview of the factors affecting midbrain organoid culture and dataset generation. iPSCs from healthy control (WT) and GBA-PD patients were derived into NESCs. For Dataset 1, midbrain organoids from early (∼10-15) or late (∼16-20) passages were generated in four independent batches. On day 30 of midbrain organoid culture, RNA sequencing and immunofluorescence experiments were performed. For Dataset 2, four independent batches of midbrain organoids were generated. On day 30 or day 60 of midbrain organoid culture, RNA sequencing and metabolomic analysis were performed.

**Table 1:**
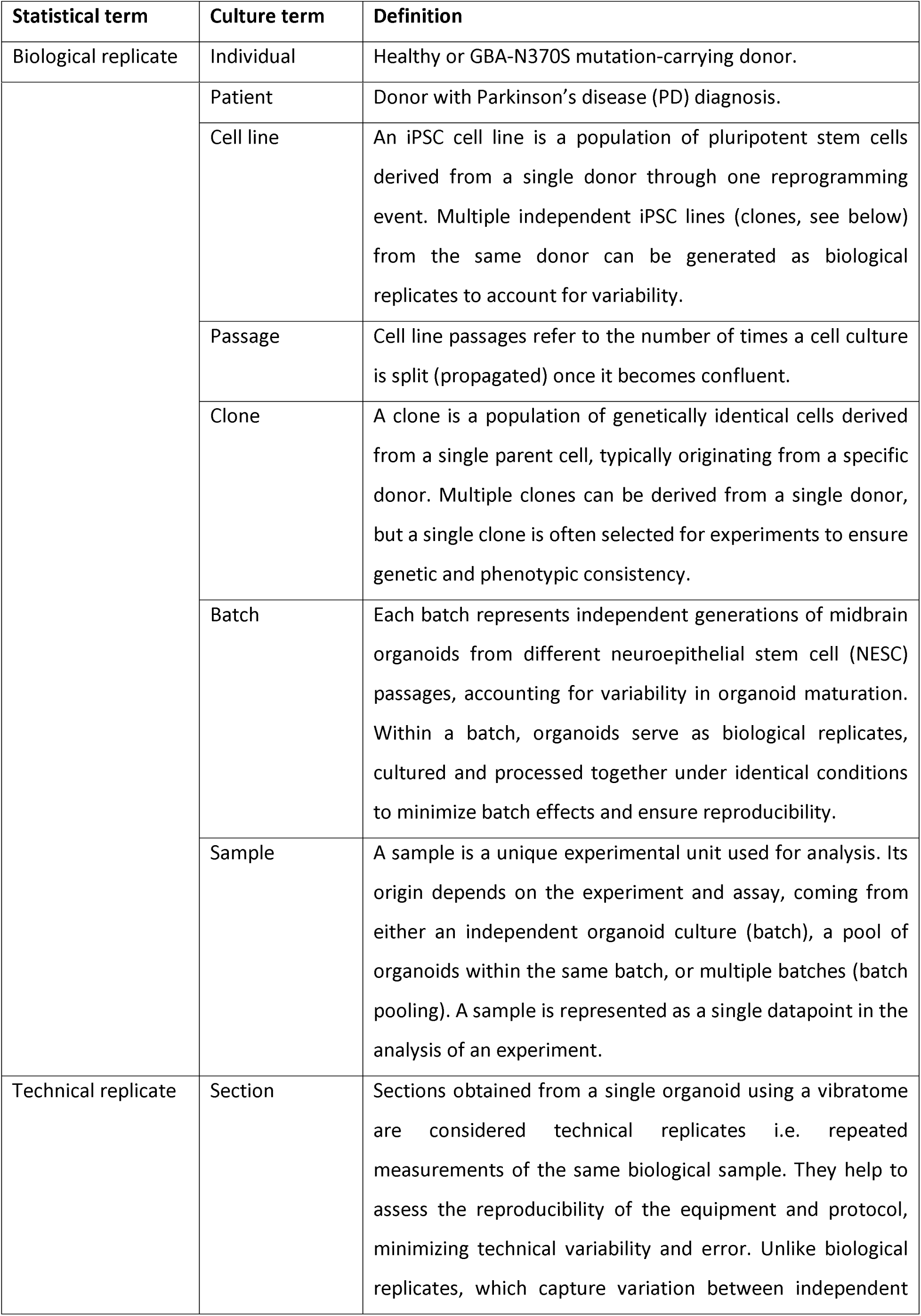
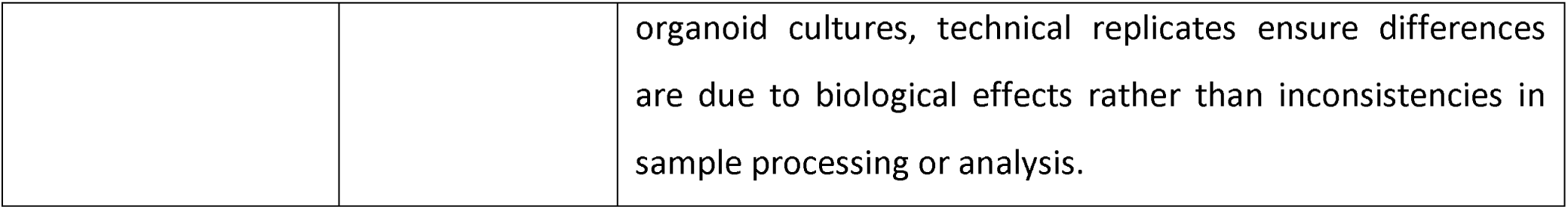
Nomenclature.

Correlation analysis of transcriptomic signatures of eight independent organoid generation batches from early and late NESC passages revealed that the features such as disease, cell line, and donor sex are interdependent factors, representing the donor (patient or healthy individual). However, the passage of NESCs and the organoid generation batch are independent sources of variance (Figure 2a). Applying Principal Variance Component Analysis (PVCA), a statistical method for quantifying and prioritizing sources of variance, revealed that the interaction between disease and sex (31.7 %) contributed most to the variance in the transcriptomic data. The passage (31%) appeared as the second highest contributor, followed by the residual variance (18.6%) which represents the variance that cannot be attributed to specific explanatory variables or known factors in our dataset. The variance attributed to batch (5%) or its interaction with passage (0.7%), sex (0%) or disease (0%) showed an insignificant role in the data variance (Figure 2b).

**Figure 2.**
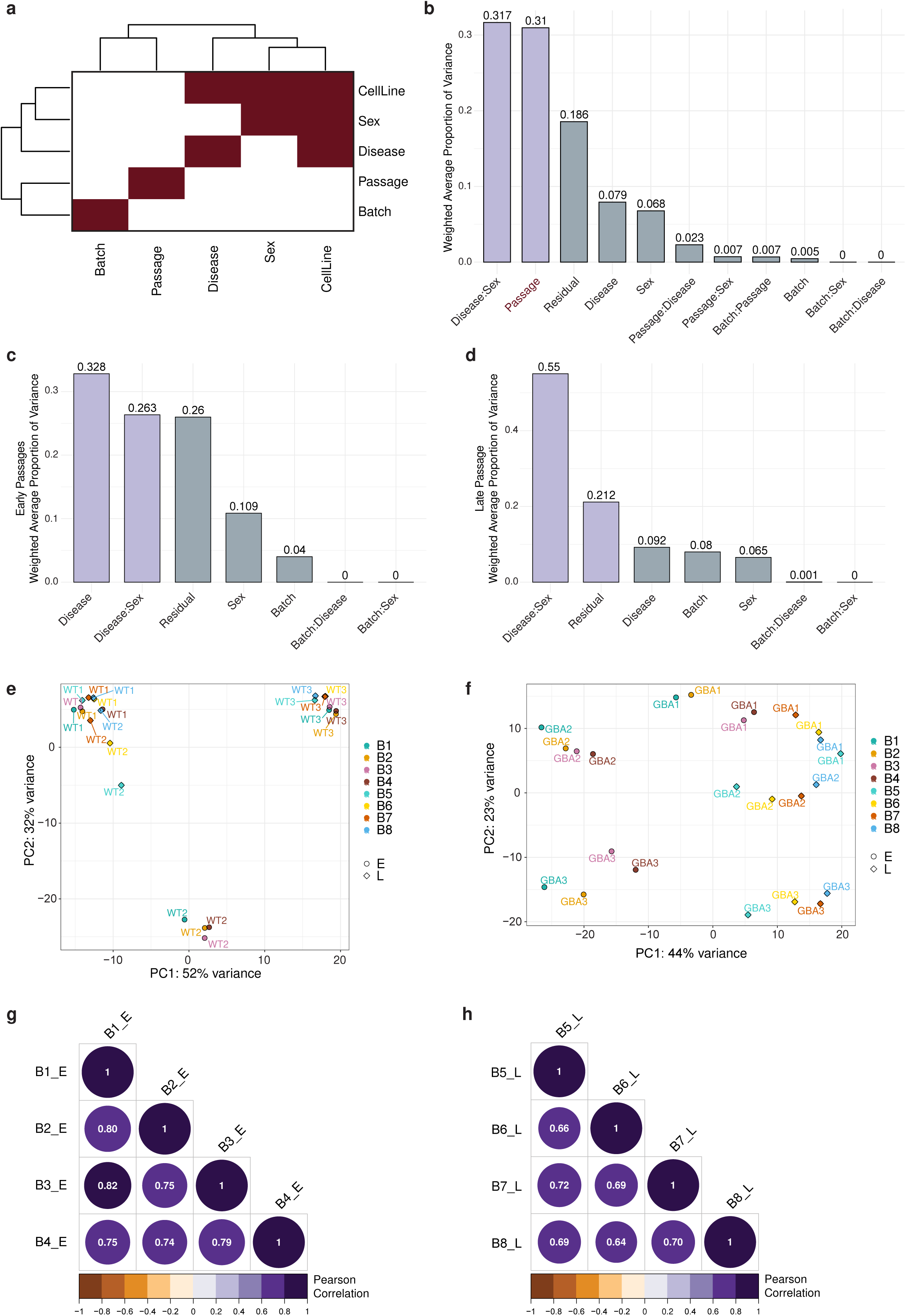
Principal Variance Component Analysis identifies the disease and the passage to be the major sources of variation in midbrain organoid culture. Midbrain organoids were generated at early or late passage in four independent batches and transcriptomic analysis was performed. (a) Correlation analysis of variance features including Batch, Passage, Disease, Sex and Cell Line. (b) Principal Variance Component Analysis (PVCA) on transcriptomic data including all passages. (c) Principal Variance Component Analysis (PVCA) on early passage transcriptomic data. (d) Principal Variance Component Analysis (PVCA) on late passage transcriptomic data. (e) Principal Component Analysis (PCA) on healthy control (WT) samples from early and late passage and all eight batches. (f) Principal Component Analysis (PCA) on PD-GBA samples from early and late passage and all eight batches. (g) Pearson Correlation of log2 fold changes (FC) of significant genes (p < 0.05) between batches derived from early passage NESCs. h) Pearson Correlation of log2 fold changes (FC) of significant genes (p < 0.05) between batches derived from late passage NESCs.

Considering the presumed role of the passage number in transcriptomic data variation, we divided the dataset in batches generated from early (pE10-15) or late NESC passages (pL16-20) to explore the effects of passage on data reproducibility in detail. The PVCA showed that the disease accounted for the largest proportion of the variance (32.8%) in organoid batches generated from early passage NESCs, followed by the interaction between disease and sex (26.3%) and the unexplained residuals (26%). The percentage of the batch was elevated compared to the complete dataset (4%) (Figure 2c). Similarly, the disease-sex interaction with 55% was the highest contributor to the variance in the organoid batches produced from the late passages (B5-8), followed by the residual variance (21.2%). We observed that the proportion of variance of the batch doubled in organoids generated from late passage NESCs compared to the early passage NESCs (8%) (Figure 2d, Supplementary Figure 1a), indicating that the higher batch-to-batch-variability is a result of later initial passages of NESCs used for organoid generation. Next, we investigated the effects of passages and batches on each cell line’s individual transcriptomic profile. Using principal component analysis (PCA), we observed that organoid samples from healthy donors (WT) tended to cluster separately, demonstrating biological variance which is independent of passage. Only in the WT2 cell line, we observed an effect of passage, with WT2 progressively resembling the WT1 cell line with each passage. This indicates that the WT2 cell line may exhibit greater variability in phenotype depending on the NESC passage (Figure 2e). Contrary, GBA-PD midbrain organoid samples clustered rather depending on the initial NESC passages, indicating that patient samples are more sensitive to experimental design and culture conditions (Figure 2f, Supplementary Figure 1b).

To explore the passage effect on a disease signature, we correlated the expression of identified significantly differentially expressed genes (DEGs) between healthy controls and GBA-PD midbrain organoids generated in batches from early or late NESC passages. Early passage batches (B1-4) showed a significant correlation of DEG expression (0.74 – 0.82) (Figure 2g), while late passage batches (B5-8) showed on average moderate correlation (0.64 – 0.72) (Figure 2h). Similarly, looking at the expression of all shared genes across the late or early passage batches, we saw a higher (0.51 – 0.64) correlation for the early passage batches than for the late passage batches (0.42 – 0.50) (Supplementary Figure 1c and d).

Overall, exploration analysis of potential sources of variation in the transcriptomic data of midbrain organoids suggests that disease status and number of passages are major sources of gene expression variance, while batch effects are minimal. Moreover, early passages show more consistent transcriptomic profiles, while late passages lead to increased variability, emphasizing the need to consider NESC passage in study design.

### Reproducible transcriptomic disease signature across independently generated datasets

The passage number showed a more considerable impact on the data reproducibility than organoid batches. We further wanted to assess more in detail how the choice of NESC passages used for organoid generation might influence the reproducibility of disease phenotypes.

We grouped the early passage batches (B1-4) and the late passage batches (B5-8) and identified the significant DEGs within these data subsets. We found 27 overlapping DEGs in the early passage set, whereas the late passage batches did not show shared DEGs (Supplementary Figure 2a and b). This could be due to the lower gene expression correlation between organoid batches from late passage NESCs, representing higher data variability. Since the DEG expression signature at early passages appeared more consistent across organoid batches, we presumed 27 DEGs shared among the early passage batches as a potential key GBA-PD disease signature and examined its reproducibility in both early and late passage batches. Here, we observed that the clustering of WT versus GBA-PD samples for the 27 DEGs was notably more distinct in the early passage heatmap compared to the late passage heatmap (Supplementary Figure 2c and d), where we still observed adequate separation. Moreover, by combining all eight batches, the expression of selected 27 DEGs also separated WT and GBA-PD samples, indicating the relevance of these genes in disease pathogenesis (Figure 3a). Enrichment analysis of the 27 significant DEGs between GBA-PD and WT samples revealed an association with Reactome pathways related to cellular senescence and apoptosis, consistent with findings of recent studies (Figure 3b) (Rosety et al., 2023). Additionally, a Gene Ontology enrichment analysis supported these results and identified “bleb assembly”, which is linked to apoptosis, as the most enriched pathway, further confirming the disease signature in GBA-PD (Figure 3c). Next, we classified the 27 genes into distinct functional categories. These included the senescence/apoptosis pathway, biological processes related to synapse and nervous system development, metabolic and oxidative stress processes, cell adhesion and extracellular matrix (ECM) dynamics, and transcription and gene regulation. Importantly, the fold change (FC) of GBA-PD vs WT demonstrated a highly similar dysregulation trend for all genes across all eight batches (Supplementary Figure 2e).

**Figure 3.**
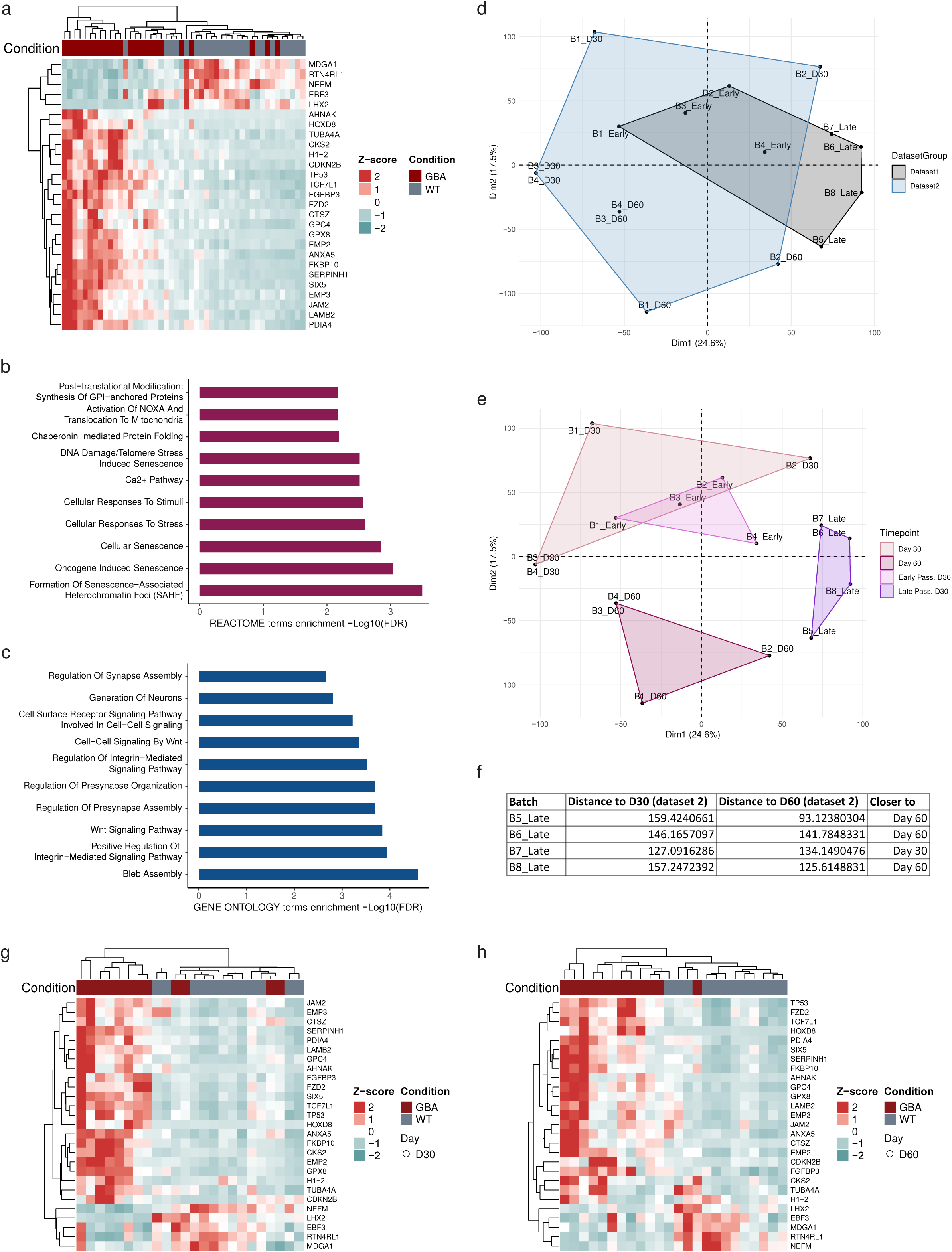
Disease signature is reproducible across independent datasets at transcriptomic level. For Dataset 1, midbrain organoids were generated at early or late passage in four independent batches and transcriptomic analysis was performed. For Dataset 2, midbrain organoids were generated in four independent batches at two timepoints and transcriptomic analysis was performed. (a) Unsupervised hierarchical clustering of GBA-PD and healthy control (WT) samples based on normalised gene counts of 27 predefined genes of all passages. (b) REACTOME pathway enrichment analysis of the DEGs between GBA-PD and healthy control (WT) samples. (c) GO pathway enrichment analysis of the DEGs between GBA-PD and healthy control (WT) samples. (d) Integration of log2 fold change (FC) from dataset 1 and dataset 2 shown in principal component analysis (PCA) plot. Samples are clustered by dataset. (e) Integration of log2 fold change (FC) from dataset 1 and dataset 2 shown in principal component analysis (PCA) plot. Samples are clustered by timepoint and passage. (f) Euclidean distance table showing the batch, the calculated distance to day 30 or day 60, and the proximity to one of the two time points. (g) Unsupervised hierarchical clustering of GBA-PD and healthy control (WT) samples based on normalised gene counts of 27 predefined genes at day 30. (h) Unsupervised hierarchical clustering of GBA-PD and healthy control (WT) samples based on normalised gene counts of 27 predefined genes at day 30.

The initial dataset included organoid transcriptomic data from a single time point - day 30 of organoid culture. We produced new independent organoid batches and performed RNA sequencing and metabolomics analyses for organoids at two-time points, day 30 and day 60 (Dataset 2). These analyses were done to additionally assess how reproducible the phenotypes are at different time points of organoid culture and across independent experiments. The transcriptomic analysis allowed us to compare the two datasets, where the PCA showed, as expected, a tighter clustering for dataset 1 (day 30 organoids) compared to a broader pattern in dataset 2 (day 30 and day 60 organoids) (Figure 3d). Early passage samples (B1-4_Early) from dataset 1 clustered with day 30 samples from dataset 2 (B1-4_D30), showing a similar expression profile (Figure 3e). In contrast, late passage samples (B5-8_Late) from dataset 1 formed a separate cluster, highlighting the significant impact of passage number on gene expression (Figure 3e). To assess the similarity of specific data points to predefined clusters, we calculated the Euclidean distance of individual points from the centroids of designated groups. The analysis revealed that most late passage samples (B1-4_Late) aligned more closely with day 60 samples (B5-8_D60), except for “B7_Late” which clustered near day 30 (B1-4_D30) (Figure 3e). These findings indicate that late passage midbrain organoids display phenotypes more similar to those observed in longer-term midbrain organoid cultures, which may reflect age-related changes in the organoid system. Next, we wanted to determine whether the samples in the newly generated dataset shared the same disease signature defined by the 27 significant DEGs common to the early passage NESC organoid batches. We observed that the WT and GBA-PD samples showed an even clearer clustering at day 60 compared to day 30. This finding aligns with expectations as day 60 midbrain organoids are more aged, leading to the accumulation of cellular, epigenetic, and transcriptional changes that enhance the distinction between sample groups (Figure 3g and h).

Altogether our analysis shows that while NESC passage can influence phenotype detection, the disease signature identified in early passages is reproducible in later passages and across independent sequencing experiments from two distinct organoid culturing time points. This establishes the midbrain organoid model as a reliable tool for disease phenotyping and modelling.

### Validation of dopaminergic neuron and senescence phenotypes at the protein level confirms transcriptomic disease signature in midbrain organoids

Our transcriptomic analysis identified a distinct disease signature in GBA-PD, characterized by significant dysregulation of genes associated with pathways related to cellular senescence and apoptosis, as well as differences in synaptic and nervous system development. To validate these transcriptomic results and confirm phenotype reproducibility at the protein level, we performed whole mount staining of midbrain organoids. We used the dopaminergic neuron marker TH (tyrosine hydroxylase), the neuronal marker MAP2 (Microtubule-associated protein 2), and the DNA damage/senescence marker 53BP1 (p53-binding protein 1) to additionally assess dopaminergic neuron and the senescence phenotypes at the protein level. We observed that GBA-PD midbrain organoids exhibit a significantly reduced number of TH-positive dopaminergic neurons compared to WT midbrain organoids consistent with results previously reported by Rosety et al (2023) and Zagare et al (2024) (Figure 4a and b). Similarly, 3D reconstruction of dopaminergic neurons showed that neurites in patient-specific organoids are shorter and less ramified compared to WT (Figure 4c). Assessing the TH transcript counts in the transcriptomic data, we observed a significant decrease in the GBA-PD cell lines, supporting the immunofluorescence findings (Supplementary Figure 3a). The log2 FC of the TH expression in GBA-PD vs. WT appeared more heterogeneous in late passage batches (B5-8) compared to the early passage batches (B1-4) (Supplementary Figure 3b), confirming that early passage batches are more reproducible at day 30 of organoid culture. Additionally, we observed that increasing the sample size through individual batches plays a critical role in detecting even the most important phenotypes (Supplementary Figure 3c and d).

**Figure 4:**
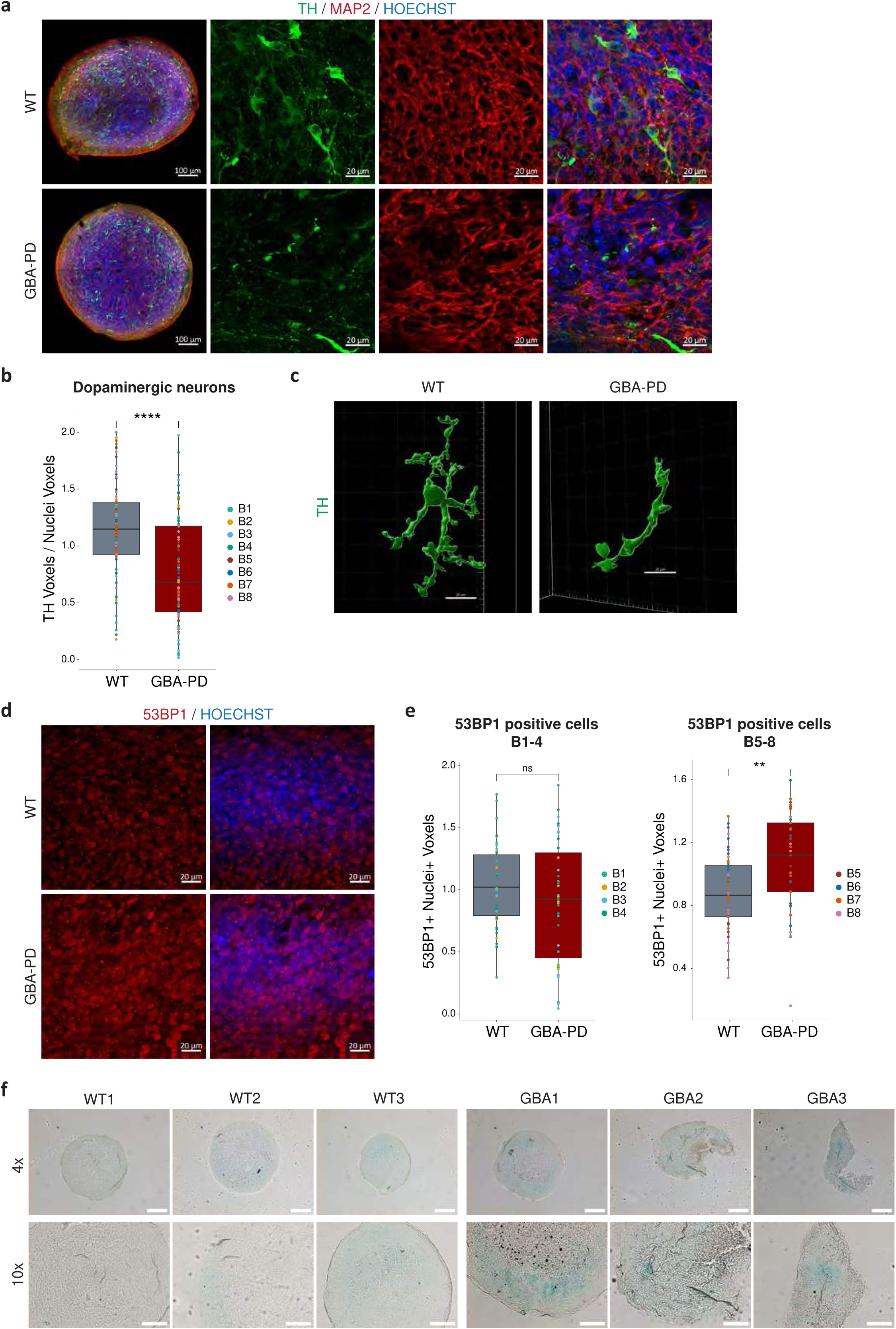
Reproducible dopaminergic neuron and senescence phenotype at protein level in midbrain organoids. Midbrain organoids were generated at early or late passage in four independent batches and whole mount immunofluorescence and β-galactosidase staining was performed. (a) Representative confocal images of TH (green), MAP2 (red) and nuclei (blue) of midbrain organoid section staining at day 30 (scale bar, 100 μm, 20x; 50 μm, 63x). (b) High-content automated image analysis of immunofluorescence stainings of TH+ dopaminergic neurons in midbrain organoids normalized by total nuclei. Data represents a summary of eight independent batches normalized to the mean of the controls per batch. Wilcoxon T-test; ****p < 0.0001. (c) 3D reconstruction with IMARIS software showing the dopaminergic neuron (TH) in the healthy control (WT) and GBA-PD cell line (scale bar = 20 μm). (d) Representative confocal images of 53BP1 (red) and nuclei (blue) in whole mount staining of midbrain organoids at day 30 (scale bar, 20 μm, 63x). (e) High-content automated image analysis of immunofluorescence stainings of 53BP1 (53BP1+) and Nuclei (Nuclei+) double-positive cells in midbrain organoids. Data represents early passage (B1-4), or late passage (B5-8) batches normalized to the mean of the controls per batch. Wilcoxon T-test; **p < 0.01. (f) Senescence-associated β-galactosidase staining (blue) of midbrain organoids (scale bar, 200 μm, 4×; 100 μm, 10×).

Analysis of the DNA damage/senescence marker 53BP1, showed that the senescence phenotype at the protein level appears in the late passage batches (B5-8), while the transcriptomic signature indicated a significant senescence-associated phenotype already in the early passage batches (B1-4) (Figure 4d and e, Supplementary Figure 4a-b). To further confirm the senescence phenotype in GBA-PD samples, we stained the organoids with β-galactosidase, a well-established marker for detecting cellular senescence (Hernandez-Segura et al., 2018). We observed a more intense β-galactosidase signal in GBA-PD midbrain organoids (Figure 4f), providing additional evidence of increased senescence in midbrain organoids derived from PD patients (Rosety et al., 2023).

### GBA-PD midbrain organoids demonstrate a reproducible metabolic profile

After confirming the phenotype reproducibility at transcriptomic and protein levels, we wanted to assess how reproducible the GBA-PD metabolic signature was. Metabolism is highly dynamic, and therefore metabolite levels may exhibit greater variability between organoid batches compared to changes in transcript or protein abundance. Nevertheless, our data showed that the majority of measured metabolite expression patterns were consistent across four batches at both time points (Figure 5a, 5b). As an exception, we identified B4 to display the most distinct metabolite abundance profile compared to the other three batches at day 30 of organoid culture. Accordingly, correlation analysis showed a significant correlation of metabolite FCs (GBA-PD vs. WT) for B1, B2 and B3 (0.61 – 0.86), while B4 demonstrated a low correlation with any of these batches (0.12) (Figure 5c). However, at the later time point (day 60), all four batches showed a significant correlation (0.59 – 0.79), suggesting that the differences in B4 at day 30 might be due to an organoid differentiation path that transiently diverges from the other batches but these differences even out in more mature organoid cultures (Figure 5d). Moreover, PCA analysis demonstrated that the metabolic profiles of WT and GBA-PD midbrain organoids are not influenced by batch effects but are instead driven by sample and disease state, further supporting the reliability and reproducibility of the data (Figure 5e).

**Figure 5:**
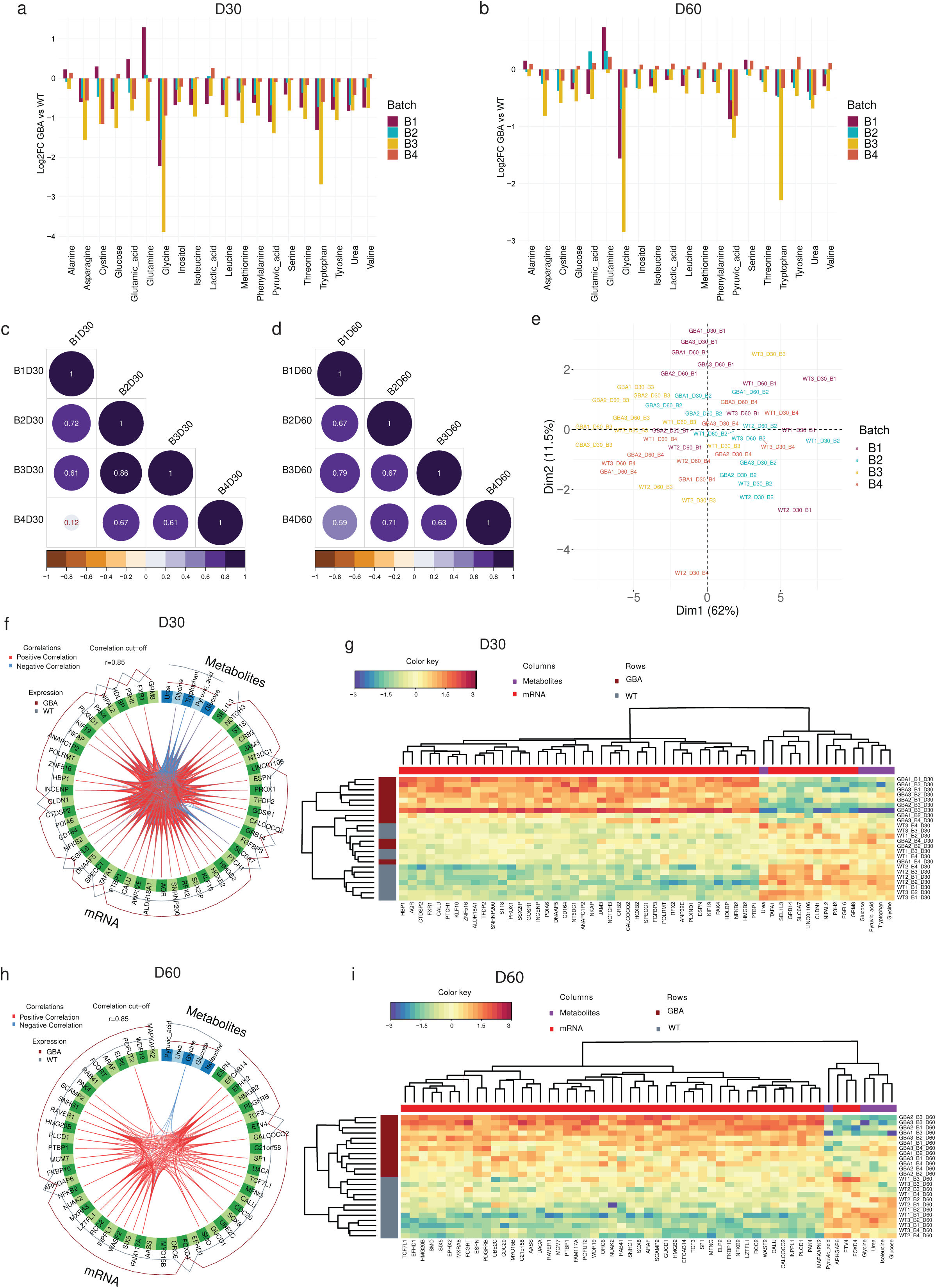
Midbrain organoids from day 30 and day 60 show a reproducible metabolic profile. (a) Metabolite abundance difference in GBA-PD samples presented as log2 fold change (FC) compared to healthy control (WT) samples across four batches at day 30 of organoid culture. (b) Metabolite abundance difference in GBA-PD samples presented as log2 fold change compared to healthy control (WT) samples across four batches at day 60 of organoid culture. (c) Pearson correlation of log2 fold changes (FC) for all metabolites across four batches at day 30 of organoid culture. (d) Pearson correlation of log2 fold changes (FC) for all metabolites across four batches at day 60 of organoid culture. (e) Principal component analysis of metabolomics data. (f) Circos plot showing the correlation between metabolomics and transcriptomics features contributing to the variation of the component 1 at day 30 of organoid culture. Correlation threshold: r=0.9. (g) Unsupervised hierarchical clustering of the top discriminant metabolomics and transcriptomics features between the GBA-PD and healthy control (WT) samples at day 30 of organoid culture. (h) Circos plot showing the correlation between metabolomics and transcriptomics features contributing to the variation of component 1 at day 60 of organoid culture. Correlation threshold: r=0.84. (i) Unsupervised hierarchical clustering of the top discriminant metabolomics and transcriptomics features between the GBA-PD and healthy control (WT) samples at day 30 of organoid culture.

In addition, we performed integrative analysis between the metabolomics and transcriptomics datasets to identify the key molecular interactions across different omics levels at different organoid culturing stages (day 30 and day 60). Importantly, this analysis confirmed earlier findings, demonstrating that the top transcripts and metabolites more effectively distinguish samples by condition rather than by batch (Supplementary Figure 5). We observed that pyruvic acid, urea, glycine, and glucose were reappearing among the top five metabolites separating the WT and GBA-PD samples at both time points (Figure 5a-i). Moreover, at both time points, the change in glycine abundance between conditions was negatively correlated with a subset of the top 50 genes, suggesting that glycine metabolism and its transcriptional regulation play a substantial role in GBA-PD (Figure 5f and h). Aligned with overall metabolism adaptations usually observed during aging and the metabolic alterations associated with GBA mutation-driven dysregulation and senescence, we found improved sample separation based on top transcriptomic and metabolomic features at day 60 of organoid culture (Figure 5g and i). This finding suggests that the metabolic phenotype of GBA-PD intensifies over time (Figure 5g and i).

Altogether these data suggest that even highly dynamic metabolic changes are reproducible in midbrain organoid models and therefore can reliably capture critical molecular interactions driving GBA-PD metabolic dysregulation.

### Sample estimation using power analysis for iPSC-derived models

A considerable limitation of studies using iPSC-derived models is the availability of cell lines, particularly when it comes to patient specific lines with a particular mutation. This leads to underpowered studies, increasing the risk of reporting false negative results (Type II error) (Brunner et al., 2023). We observed exactly this in our here presented example study on TH gene expression, where a single batch with three samples per group failed to capture the true differences in TH expression levels between the control and GBA-PD samples. Therefore, using independent experimental batches can be used to enhance the experimental throughput by increasing sample sizes. Thus, in addition, to assessing the reproducibility of the midbrain organoid model, we wanted to estimate the required sample size for RNA sequencing, metabolomics and imaging experiments to reach an optimal statistical power of 80%, considering the presented data as pilot experiments for future experimental designs using organoid models. 80% statistical power is by convention the target power of a study to reduce the probability of rejecting a false null hypothesis (Serdar et al., 2020). For RNA sequencing datasets, we estimated how many samples per group are required to reach optimal statistical power based on the observed effect size as log2 FC between the significant DEGs in this study. While the absolute values of log2 FC ranged from 0.08 (min) and 8.21 (max), the median was 0.42, indicating that for most of the DEGs the difference between the two groups is relatively small and 30 samples per group would be required to detect true differences in the gene expression (Figure 6a and d). This means that for the most differentially expressed transcripts, 10 batches must be generated to detect true significant differences between the groups if we have 3 cell lines from each group or condition available. For a larger effect size between groups (>1.5 log2 FC), the sample size required for optimal power of 80% decreases to 9 samples per group. Consequently, three independent rounds of organoid generation are required to detect significantly differentially expressed genes with an expected expression difference greater than 1.5-fold when three cell lines per group or condition are used. In the metabolomics dataset, we estimated the effect size as Cohen’s d of each significantly differentially abundant metabolite. Median d was large 0.91, which would require 20 samples per sample group to reach optimal statistical power in the analysis (Figure 6b and d). The maximum d was 1.58, decreasing the required samples number to ten, while the smallest d was 0.73, increasing the required sample number to 35 per group (Figure 6b and d). In the imaging dataset, we estimated the required sample size based on the values of TH normalised by nuclei, as this is the expected imaging feature to be significantly different between the healthy control and GBA-PD samples. With the calculated effect size d=0.56, 50 samples per group would be necessary to reach 80% statistical power (Figure 6c and d). In organoid studies, immunofluorescence stainings are done on organoid sections. Unlike RNA sequencing or metabolomics experiments, where organoids are pooled into a single sample represented by one data point in the analysis, in imaging experiments we consider each individual organoid as a biological replicate. This increases the sample size, which allows for optimal statistical power considering the rather low expected effect size between the sample groups in imaging.

**Figure 6.**
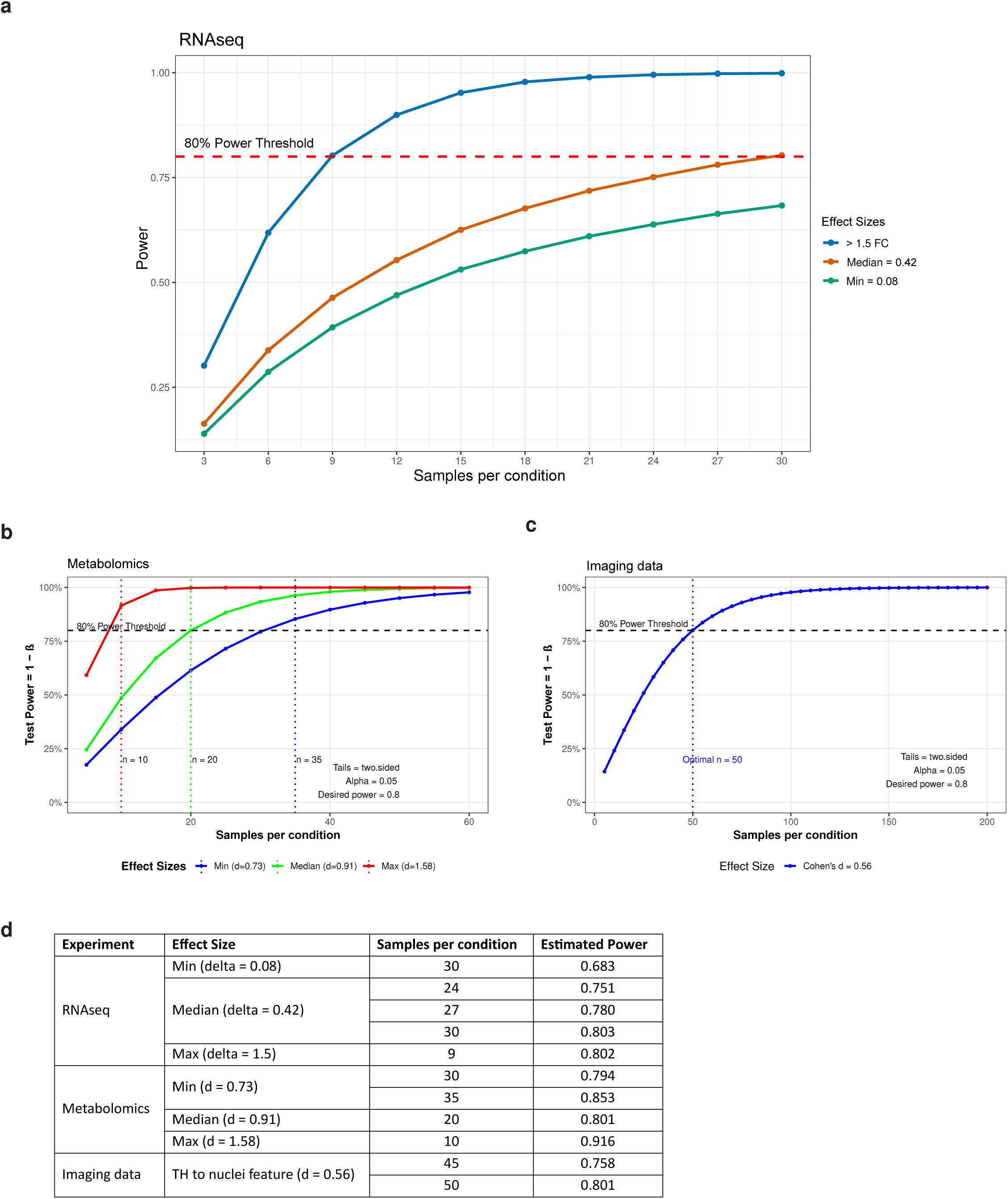
Sample Estimation Using Power Analysis for three data types (RNAseq, Metabolomics, Imaging). Each panel plots statistical power (1 – β) against sample size per group at α = 0.05, highlighting the sample sizes needed to reach the 80% power threshold (dashed line) for different effect sizes determined from this pilot study in each data type. (a) RNAseq shows effect sizes of >1.5-fold (blue), median = 0.42 (orange), and min = 0.08 (green). (b) Metabolomics compares min (d = 0.73), median (d = 0.91), and max (d = 1.58) effect sizes; vertical lines mark the number of replicates at which each effect achieves 80% power. (c) Imaging data illustrates a single effect size (Cohen’s d = 0.56), requiring ∼n = 50 per group to reach 80% power. (d) Table summarising the power analysis results showing the effect size, the sample size per group and the estimated power for each experiment performed in this study (RNAseq, metabolomics and imaging data).

For instance, in this pilot study the use of three cell lines and eight batches per group increases the sample size to 24 per group. Similar approaches can be used to achieve the optimal sample size, considering the specific study objective and the standards required (Jensen et al, 2023).

This analysis highlights the importance of incorporating multiple batches to achieve statistically robust and reproducible results, when the availability of biological replicates (cell lines) is limited. However, increasing the number of cell lines is always preferred over adding more batches, as it results in more generalizable and translatable findings.

## Discussion

Our study establishes midbrain organoids as a reproducible in vitro model to study Parkinson’s disease. By generating multiple independent organoid batches and examining both early and late passages, as well as different time points, we address a key challenge in the field: ensuring reproducibility and consistency in complex 3D culture systems.

Our findings reveal that, despite the inherent complexity of organoid models, batch effects can be successfully minimized. The principal variance component analysis underscores that donor-related factors, disease state, and sex, are prominent drivers of transcriptional variability, while passage of the NESCs emerges as a technical parameter influencing transcriptomic stability. Notably, early-passage cultures consistently yield a more robust and stable disease-specific transcriptomic signature compared to late-passage organoids, which nonetheless still maintain core disease-related features. This signature, associated with TH-positive dopaminergic neurons loss and cellular senescence, aligns with previously reported phenotypes in GBA-PD models and reflects key pathogenic mechanisms implicated in disease progression (Rosety et al., 2023; Zagare et al., 2024).

Crucially, we validated these transcriptomic differences at the protein and metabolic levels. Changes in dopaminergic neuron numbers, along with increased senescence-associated markers, were consistent with our gene expression data. Furthermore, the metabolic profiles were largely stable and reproducible across independent batches of organoid generation, particularly as the organoids matured. This evidence supports the notion that midbrain organoids can recapitulate key aspects of PD pathology reproducibly along multi-level analysis.

Although this study focused on PD patient midbrain organoid samples carrying the GBA-N370S genetic variant, we believe that the number of samples from independent organoid generation batches was sufficient to assess the overall variation in the organoid data. Thus, our conclusions on organoid reproducibility can likely be generalized to other organoid experiments involving two or more independent sample groups. Furthermore, our analysis incorporated sample size estimation to achieve optimal statistical power across bulk RNAseq, immunofluorescence high-content imaging, and metabolomics experiments, offering guidelines for designing future organoid studies to ensure result accuracy and minimize false positive results. It is important to note that midbrain organoids represent a guided brain organoid model, where variability between individual organoids and batches is expected to be lower compared to unguided brain organoid protocols. In such unguided models, larger sample sizes may be necessary to reliably detect true differences between sample groups.

While including batches can enhance the statistical power of a study, they should not be used to artificially inflate power. Instead, batches should serve as an additional replicate alongside multiple distinct cell lines (if available) or used as a strategy to help overcome the common challenge of limited cell line availability (e.g. in the case of rare mutations).

In conclusion, this study provides strong evidence that patient-derived midbrain organoids are not only an accurate model of GBA-PD pathology but are also a robust and reproducible experimental system. By highlighting and controlling critical sources of variability, these models can build a solid foundation for future work aimed at unravelling disease mechanisms and accelerating the development of personalized treatments for Parkinson’s disease.

## Supporting information

RNA-Seq Metadata

## Data availability

All original and processed data as well as scripts that support the findings of this study are public available at this https://doi.org/10.17881/4n7e-vc7. Gene expression datasets can be accessed on Gene Expression Omnibus (GEO) under the accession codes GSE287566 and GSE269316.

## Code availability

All scripts used to obtain, analyse and plot the data are available at https://gitlab.com/uniluxembourg/lcsb/developmental-and-cellular-biology/zuccoli_2025.

## Acknowledgments

We thank the LCSB Metabolomics and Lipidomics Platform for their service. We acknowledge Novogene for RNA sequencing. We would like to thank the LCSB Bioimaging Platform for supporting high-content imaging and image analysis workflow. We would also like to thank Dr. Sònia Sabaté Soler for the help with the 3D reconstruction using the IMARIS software and Dr. Matthieu Gobin for his critical contributions to the manuscript.

This project has received funding from the National Centre of Excellence in Research on Parkinson’s Disease (NCER-PD) which is funded by the Luxembourg National Research Fund (FNR) (FNR/NCER13/BM/11264123). Further, FNR funding as part of the i2TRON Doctoral Training Unit (PRIDE19/14254520) and the CORE program (CORE 22_BM_17193204_MidStriPD) is acknowledged.

## Rights retention statement

This research was funded in whole by the FNR-Luxembourg. For the purpose of Open Access, the author has applied a CC BY public copyright license to any Author Accepted Manuscript (AAM) version arising from this submission.

## Author Contributions

E.Z. and A.Z. conceived, designed and collected data. E.Z., H.A. and A.Z. performed data analysis and interpretation of results. M.T. contributed with experiments. The work was supervised by J.C.S., E.Z., H.A. and A.Z. wrote the original manuscript. S.N. and J.C.S. revised and edited the manuscript.

## Figure Legends

**Supplementary Figure 1: Correlation analysis of healthy control (WT) and GBA-PD samples.**

(a) Unsupervised hierarchical clustering plot showing the correlation of healthy control (WT) samples from early and late passage and all eight batches. (b) Unsupervised hierarchical clustering plot showing the correlation of GBA-PD samples from early and late passage and all eight batches. (c) Pearson Correlation of log2 fold changes (FC) of all genes between batches derived from early passage NESCs. (d) Pearson Correlation of log2 fold changes (FC) of all genes between batches derived from late passage NESCs.

**Supplementary Figure 2: Identification of 27 differentially expressed genes (DEGs) in early passage batches and assignment to distinct functional categories.**

(a) Venn diagram of DEGs showing the batches derived from early passage NESCs, with 27 genes in common. (b) Venn diagram of DEGs showing the batches derived from early passage NESCs. (c) Unsupervised hierarchical clustering of GBA-PD and healthy control (WT) samples based on normalised gene counts of 27 predefined genes for early passage batches. (d) Unsupervised hierarchical clustering of GBA-PD and healthy control (WT) samples based on normalised gene counts of 27 predefined genes for late passage batches. (e) Log2 fold changes (FC) of 27 predefined genes divided into distinct functional categories. The genes were assigned to the senescence/apoptosis pathway, synapse and nervous system development, metabolic and oxidative stress processes, cell adhesion and extracellular matrix (ECM) dynamics, and transcription and gene regulation. Each category shows the FC per batch with eight batches in total.

**Supplementary Figure 3: Transcriptomic analysis of dopaminergic neuron phenotype supports imaging analysis.**

(a) TH counts from all eight batches pooled. Wilcoxon T-test; ****p < 0.0001. (b) Log2 fold change (FC) of TH expressed in early passage batches (B1-4) and late passage batches (B5-8). (c) TH counts from early passage batches (B1-4). Pooled healthy control (WT) or GBA-PD lines are shown per batch. (d) TH counts from late passage batches (B5-8). Pooled healthy control (WT) or GBA-PD lines are shown per batch.

**Supplementary Figure 4: Transcriptomic analysis of senescence phenotype supports imaging analysis.**

(a) TP53 counts from early passage batches (B1-4) pooled. Wilcoxon T-test; ****p < 0.0001. (b) TP53 counts from late passage batches (B5-8) pooled. Wilcoxon T-test; ***p < 0.001.

**Supplementary Figure 5: Partial Least Squares Discriminant Analysis**

(PLS-DA) (a) Day 30 sample discrimination by batch. (b) Day 30 sample discrimination by condition. (c) Day 60 sample discrimination by batch. (d) Day 60 sample discrimination by condition.

## References

Kalia, L. V., & Lang, A. E. (2015). Parkinson’s disease. The Lancet, 386(9996), 896–912.

Kelava, I., & Lancaster, M. A. (2016). Dishing out mini-brains: Current progress and future prospects in brain organoid research. Developmental Biology, 420(2), 199–209.

Dovonou, A., Bolduc, C., Soto Linan, V., Gora, C., Peralta III, M. R., & Lévesque, M. (2023). Animal models of Parkinson’s disease: Bridging the gap between disease hallmarks and research questions. Translational Neurodegeneration, 12(1), 36.

Monzel, A. S., Smits, L. M., Hemmer, K., Hachi, S., Moreno, E. L., van Wuellen, T., … & Schwamborn, J. C. (2017). Derivation of human midbrain-specific organoids from neuroepithelial stem cells. Stem cell reports, 8(5), 1144–1154.

Nickels, S. L., Modamio, J., Mendes-Pinheiro, B., Monzel, A. S., Betsou, F., & Schwamborn, J. C. (2020). Reproducible generation of human midbrain organoids for in vitro modeling of Parkinson’s disease. Stem Cell Research, 46, 101870.

Smith, L., & Schapira, A. H. V. (2022). GBA variants and Parkinson disease: Mechanisms and treatments. Cells, 11(8), 1261.

Chinta, S. J., & Andersen, J. K. (2018). Redox imbalance in Parkinson’s disease. Biochimica et Biophysica Acta (BBA) - Molecular Basis of Disease, 1864(2), 1078–1083.

Melo dos Santos, L. S., Trombetta-Lima, M., Eggen, B. J. L., & Demaria, M. (2024). Cellular senescence in brain aging and neurodegeneration. Ageing Research Reviews, 93, 102141.

Wang, Y., Kuca, K., You, L., &, et al. (2024). The role of cellular senescence in neurodegenerative diseases. Archives of Toxicology, 98(10), 2393–2408.

Rosety, I., Zagare, A., Saraiva, C., Nickels, S., Antony, P., Almeida, C., … & Schwamborn, J. C. (2023). Impaired neuron differentiation in GBA-associated Parkinson’s disease is linked to cell cycle defects in organoids. npj Parkinson’s Disease, 9(1), 166.

Reinhardt, P., Glatza, M., Hemmer, K., Tsytsyura, Y., Thiel, C. S., Höing, S., … & Sterneckert, J. (2013). Derivation and expansion using only small molecules of human neural progenitors for neurodegenerative disease modeling. PloS one, 8(3), e59252.

Bolognin, S., Fossépré, M., Qing, X., Jarazo, J., Ščančar, J., Moreno, E. L., … & Schwamborn, J. C. (2019). 3D cultures of Parkinson’s disease-specific dopaminergic neurons for high content phenotyping and drug testing. Advanced Science, 6(1), 1800927.

Hiltemann, S., Rasche, H., Gladman, S., Hotz, H. R., Larivière, D., Blankenberg, D., … & Batut, B. (2023). Galaxy Training: A powerful framework for teaching!. PLoS computational biology, 19(1), e1010752.

Doyle, M., Phipson, B., & Dashnow, H. (2024). 1: RNA-Seq reads to counts.

Love, M. I., Huber, W., & Anders, S. (2014). Moderated estimation of fold change and dispersion for RNA-seq data with DESeq2. Genome biology, 15, 1–21.

Benjamini, Y., & Hochberg, Y. (1995). Controlling the false discovery rate: a practical and powerful approach to multiple testing. Journal of the Royal statistical society: series B (Methodological), 57(1), 289–300.

Kuleshov, M. V., Jones, M. R., Rouillard, A. D., Fernandez, N. F., Duan, Q., Wang, Z., … & Ma’ayan, A. (2016). Enrichr: a comprehensive gene set enrichment analysis web server 2016 update. Nucleic acids research, 44(W1), W90–W97.

Milacic, M., Beavers, D., Conley, P., Gong, C., Gillespie, M., Griss, J., … & D’Eustachio, P. (2024). The reactome pathway knowledgebase 2024. Nucleic acids research, 52(D1), D672–D678.

Ashburner, M., Ball, C. A., Blake, J. A., Botstein, D., Butler, H., Cherry, J. M., … & Sherlock, G. (2000). Gene ontology: tool for the unification of biology. Nature genetics, 25(1), 25–29.

Aleksander, S. A., Balhoff, J., Carbon, S., Cherry, J. M., Drabkin, H. J., Ebert, D., … & Zarowiecki, M. (2023). The gene ontology knowledgebase in 2023. Genetics, 224(1), iyad031.

Shirkhorshidi, A. S., Aghabozorgi, S., & Wah, T. Y. (2015). A comparison study on similarity and dissimilarity measures in clustering continuous data. PloS one, 10(12), e0144059.

Kim, T., Chen, I. R., Lin, Y., Wang, A. Y. Y., Yang, J. Y. H., & Yang, P. (2019). Impact of similarity metrics on single-cell RNA-seq data clustering. Briefings in bioinformatics, 20(6), 2316–2326.

Bushel, P. (2024). PVCA: Principal Variance Component Analysis (PVCA) [R package version 1.46.0]. Bioconductor.

Rohart, F., Gautier, B., Singh, A., & Lê Cao, K. A. (2017). mixOmics: An R package for ‘omics feature selection and multiple data integration. PLoS computational biology, 13(11), e1005752.

Singh, A., Shannon, C. P., Gautier, B., Rohart, F., Vacher, M., Tebbutt, S. J., & Lê Cao, K. A. (2019). DIABLO: an integrative approach for identifying key molecular drivers from multi-omics assays. Bioinformatics, 35(17), 3055–3062.

Zagare, A., Hemedan, A., Almeida, C., Frangenberg, D., Gomez-Giro, G., Antony, P., … & Schwamborn, J. C. (2024). Insulin Resistance Is a Modifying Factor for Parkinson’s Disease. Movement Disorders.

Hernandez-Segura, A., Nehme, J., & Demaria, M. (2018). Hallmarks of cellular senescence. Trends in cell biology, 28(6), 436–453.

Brunner, J. W., Lammertse, H. C. A., van Berkel, A. A., &, et al. (2023). Power and optimal study design in iPSC-based brain disease modelling. Molecular Psychiatry, 28(8), 1545–1556

Smits, L. M., Reinhardt, L., Reinhardt, P., &, et al. (2019). Modeling Parkinson’s disease in midbrain-like organoids. npj Parkinson’s Disease, 5(1), 5.

Serdar, C. C., Cihan, M., Yücel, D., & Serdar, M. A. (2020, December 15). Sample size, power and effect size revisited: Simplified and practical approaches in pre-clinical, clinical and laboratory studies. Biochemia Medica, 31(1), 010502.

Jensen, K. B., & Little, M. H. (2023). Organoids are not organs: Sources of variation and misinformation in organoid biology. Stem Cell Reports.

