## Supplementary Figures for "Reproducibility of PD patient-specific midbrain organoid data for in vitro disease modelling"

**Supplementary Figure 1**

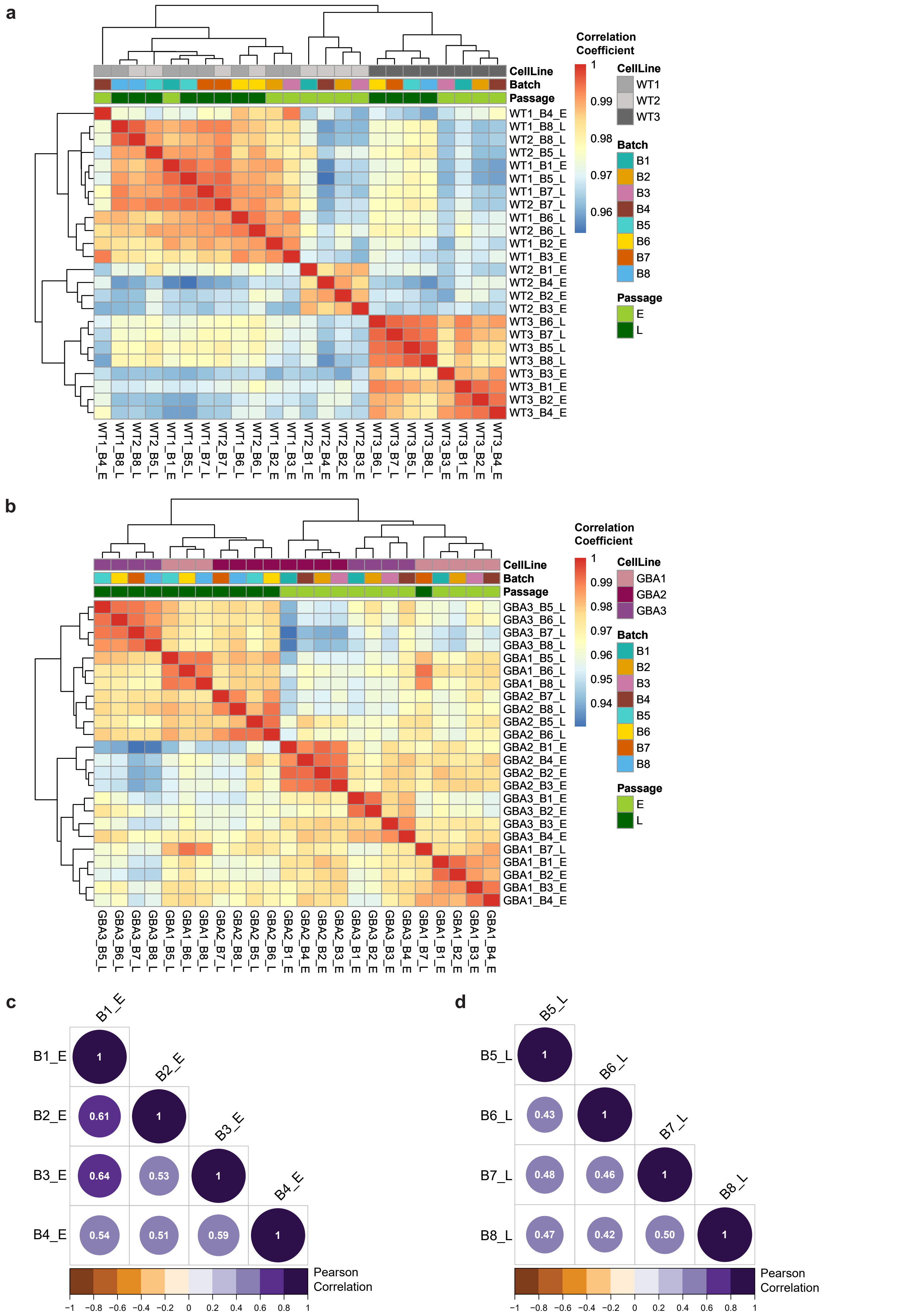

### Supplementary Figure 2

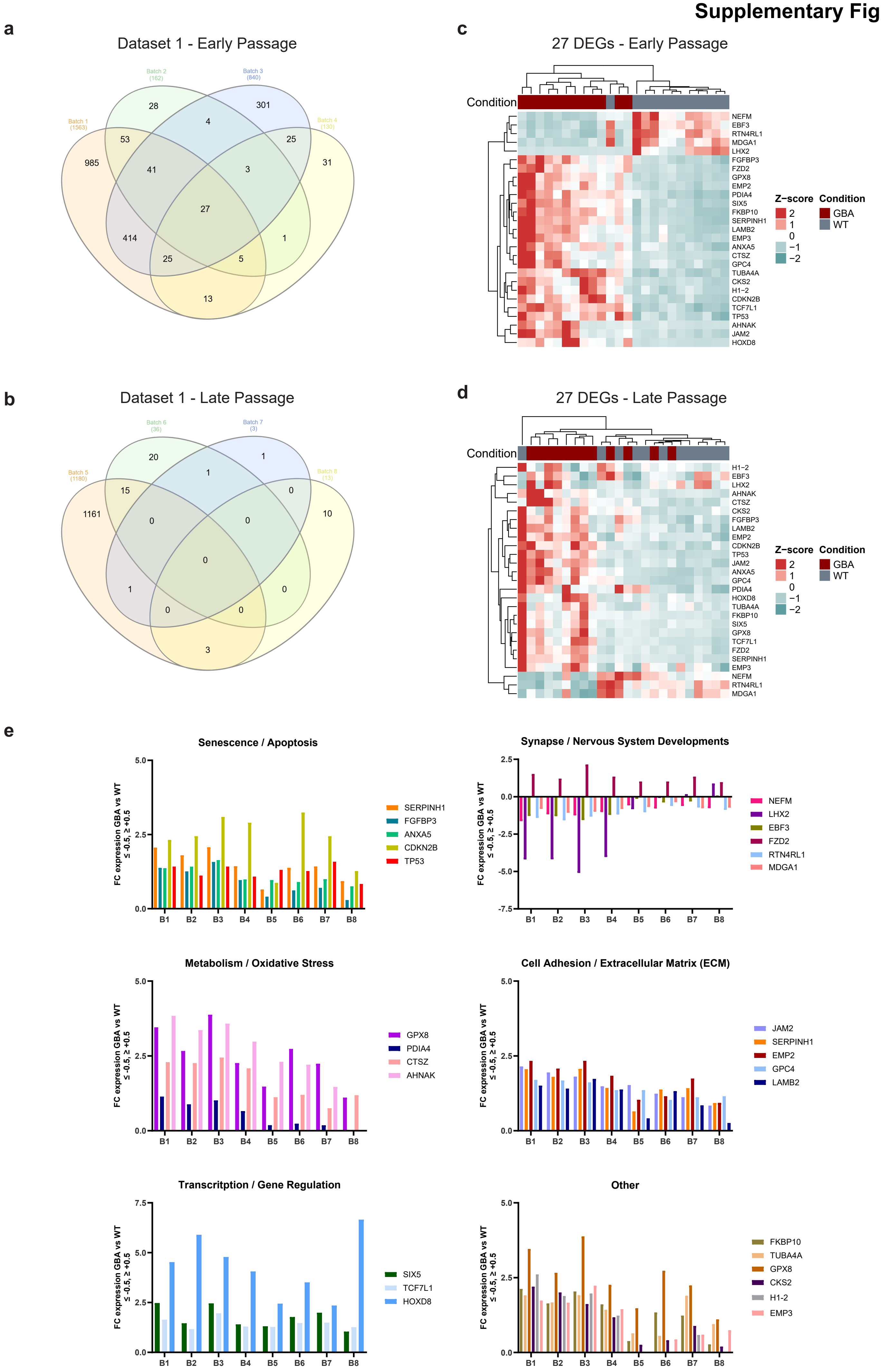

**Supplementary Figure 3**

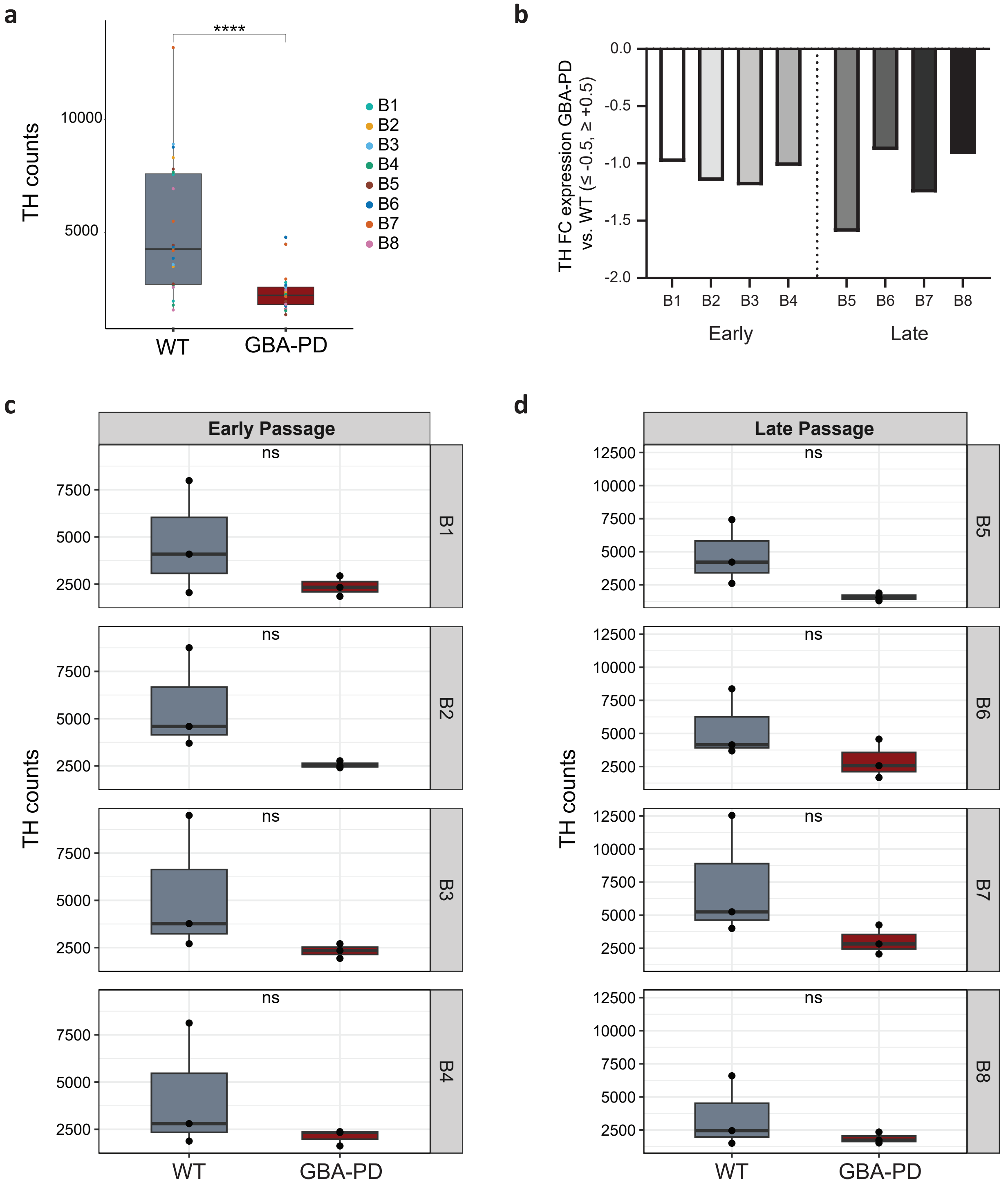

#### Supplementary Figure 4

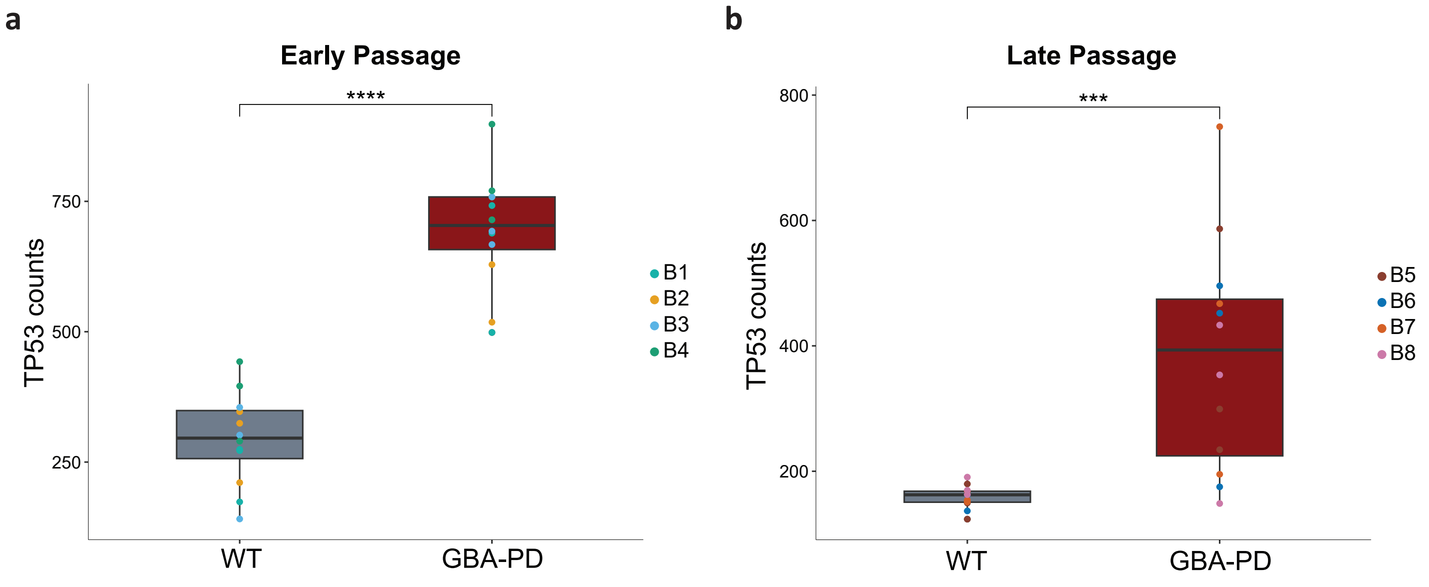

### Supplementary Figure 5

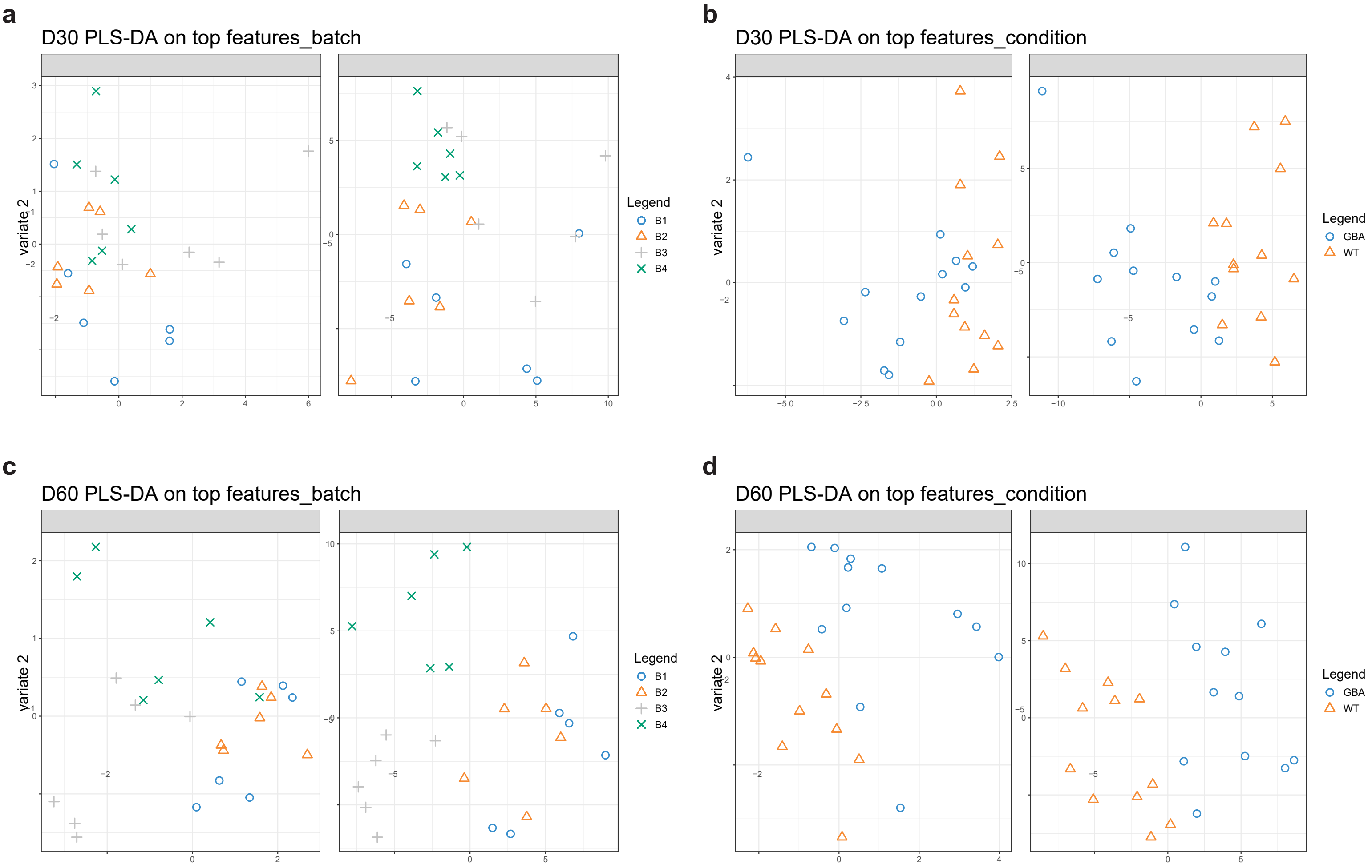
