## Supplementary Table 1 for "Reproducibility of PD patient-specific midbrain organoid data for in vitro disease modelling"

| Sample ID | Diagnosis | Genotype | Age of sampling | Source of iPSC |
| --- | --- | --- | --- | --- |
| WT1 | Healthy | wt/wt | 63 | IBBL / Max Planck Institute |
| WT2 | Healthy | wt/wt | 68 | IBBL / Max Planck Institute |
| WT3 | Healthy | wt/wt | 55 | Coriell Institute |
| GBA1 | PD | GBA-N370S/wt | 81 | European Bank for induced pluripotent stem cells |
| GBA2 | PD | GBA-N370S/wt | 55 | University College London |
| GBA3 | PD | GBA-N370S/wt | 66 | Coriell Institute |

**Table S1. Cell lines used in this study**
