## Supplementary Table 2 for "Reproducibility of PD patient-specific midbrain organoid data for in vitro disease modelling"

| Antibody | Source | Cat.no. | RRID | Species | Dilution |
| --- | --- | --- | --- | --- | --- |
| TH | Abcam | ab76442 | *AB_1524535* | Chicken | 1:250  1:1000 |
| TUJ1 | BioLegend | 801201 | *AB_2313773* | Mouse | 1:300 |
| 53BP1 | Novus | NB100-304 | *AB_10003037* | Rabbit | 1:250 |
| MAP2 | Abcam | ab92434 | *AB_2138147* | Chicken | 1:250  1:1000 |
| Anti-chicken 488 | Jackson Immunoresearch | 703-545-155 | *AB_2340375* | Donkey | 1:1000 |
| Anti-chicken 647 | Millipore | AP194SA6 | *AB_2650475* | Donkey | 1:1000 |
| Anti-rabbit 488 | Invitrogen | A21206 | *AB_2535792* | Donkey | 1:1000 |
| Anti-rabbit 568 | Thermo Fisher Scientific | A-10042 | *AB_2534017* | Donkey | 1:1000 |
| Anti-mouse 647 | Thermo Fisher Scientific | A-31571 | *AB_162542* | Donkey | 1:1000 |

**Table S2. Primary and secondary antibodies used for immunofluorescence stainings**
